# iComBat: An Incremental Framework for Batch Effect Correction in DNA Methylation Array Data

**DOI:** 10.1101/2025.05.06.652337

**Authors:** Yui Tomo, Ryo Nakaki

**Affiliations:** Japan Institute for Health Security; Rhelixa, Inc.

**Keywords:** ComBat, Empirical Bayes estimation, Epigenetics, Microarray, Repeated measurement

## Abstract

DNA methylation is associated with various diseases and aging; thus, longitudinal and repeated assessments of methylation patterns are crucial for revealing the mechanisms of disease onset and identifying factors associated with aging. The presence of batch effects influences the analysis of DNA methylation array data. Since existing methods for correcting batch effects are designed to correct all samples simultaneously, when data are incrementally measured and included, the correction of newly added data affects previous data. In this study, we propose an incremental framework for batch-effect correction based on ComBat, a location/scale adjustment approach using a Bayesian hierarchical model, and empirical Bayes estimation. Using numerical experiments and application to actual data, we demonstrate that the proposed method can correct newly included data without re-correcting the old data. The proposed method is expected to be useful for studies involving repeated measurements of DNA methylation, such as clinical trials of anti-aging interventions.

## 1 Introduction

DNA methylation is an epigenetic modification characterized by the addition of a methyl group to the cytosine base constituting a DNA molecule [1]. While DNA methylation does not modify the DNA sequence itself, it plays a vital role in the regulation of gene expression and is related to a variety of biological processes. For example, DNA methylation is associated with the onset of various diseases, from cancer to infectious diseases [2, 3, 4]. Furthermore, studies have revealed a relationship between DNA methylation and aging [5, 6, 7]. Therefore, a comprehensive analysis of DNA methylation patterns is essential for revealing the mechanisms underlying disease pathogenesis and aging.

Recently, DNA methylation arrays have been widely used to analyze large-scale samples. Epigenome-wide association studies (EWAS) have assessed and identified associations between methylation at each measured methylation site and specific phenotypes [8]. In addition, formulas known as epigenetic clocks, which calculate the biological age from DNA methylation data, have been developed using statistical and machine learning approaches [9, 10, 11]. These epigenetic clocks are currently employed to evaluate antiaging interventions and to assess the effects of aging-related exposure [12, 13]. However, batch effects present a major challenge in analyzing DNA methylation array data [14]. Batch effects are systematic variations arising from technical factors such as differences in instrumentation, reagent lots, measurement times, and other experimental conditions across batches. These effects may impede data analysis and potentially influence biological interpretations and clinical decision-making [15, 16]. Therefore, the correction of batch effects is critical for enhancing the reliability of data analyses.

Various statistical methods have been developed for batch-effect correction. Quantile normalization standardizes the distribution of signal intensities among samples under the assumption that signals of the same rank share the same intensity [17]. Surrogate variable analysis (SVA) and two-step removal of unwanted variation (RUV-2) methods adjust for unobserved sources of variability by extracting latent variables through a low-rank approximation of the residual matrix, which is obtained by regressing the signal intensity matrix on observed factors, and subsequently removing the associated variation through additional regression [18, 19]. ComBat is based on a location/scale (L/S) adjustment model that corrects data across batches by adjusting the mean and scale parameters of the observed data. In ComBat, the location and scale parameters for each gene are estimated using empirical Bayes methods within a hierarchical model that borrows information across genes in each batch [20]. ComBat has been widely adopted and extended in many studies as it works well even when sample sizes within batches are small [21, 22, 23].

In addition to post-hoc statistical corrections, preprocessing pipelines, such as SeSAMe, have been employed to reduce batch effects at the signal normalization stage. SeSAMe [24] is particularly effective in addressing technical biases stemming from array-specific features, including dye bias, background noise, and scanner variability. However, SeSAMe alone cannot fully correct all sources of batch effects. It remains limited in mitigating biological and experimental variations, such as differences in DNA extraction protocols, plate layouts, and bisulfite conversion efficiencies. These uncorrected sources of variation may affect the accuracy of the downstream DNA methylation Beta-value or M-value calculation. These findings highlighted the need for complementary statistical correction methods.

Conventional methods for batch-effect correction have been designed to simultaneously correct data from all batches. However, in long-term studies, where data are repeatedly measured and evaluated, new batches are continuously added. In such scenarios, it is desirable to apply a consistent correction to the newly included data without modifying the already corrected existing data. In this study, we developed an incremental framework for batch-effect correction based on the L/S adjustment model and the Bayesian framework employed in ComBat. The proposed method is expected to facilitate the analysis of repeatedly included new batches without recording previously corrected data and allow us to consistently interpret the results from the overall dataset.

The remainder of this paper is organized as follows: In Section 2, we describe the details of ComBat. In Section 3, we propose an incremental batch effect correction framework based on ComBat. In Section 4, we perform numerical experiments to evaluate the performance of our proposed method. In Section 5, we apply the proposed method to an actual dataset for illustrating the practical performance. In Section 6, we discuss the advantages of our proposed method and some potential directions for future works.

## 2 Batch Effect Correction Using ComBat

ComBat employs a statistical model that accounts for additive and multiplicative batch effects to eliminate these effects [20]. Furthermore, the model was designed as a Bayesian hierarchical model to borrow information across methylation sites within each batch, which is expected to perform stably even with small sample sizes. This estimation is based on an empirical Bayes approach [25].

To formulate the model and estimation method, we introduce the following notation. Let *Y*_*ijg*_ ∈ ℝ denote the M-value for batch *i*, sample *j*, and methylation site *g* (*i* = 1, …, *m, j* = 1, …, *n*_*i*_, and *g* = 1, …, *G*). The M-value is defined as the log-ratio of the intensities of the methylated and unmethylated signals:

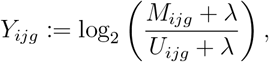

where *M*_*ijg*_ *>* 0 denotes methylated signal intensities and *U*_*ijg*_ *>* 0 denotes unmethylated [26]. Here, *λ >* 0 is a small positive constant added to numerical stability. The M-value can be approximately converted to the beta value, defined as

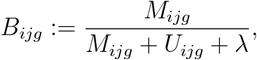

via the following transformation:

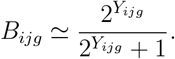

Let *X*_*ij*_ ∈ ℝ^*p*^ denote the covariate vector representing the sample conditions and let *β*_*g*_ ∈ ℝ^*p*^ denote the corresponding regression coefficients. Furthermore, let *α*_*g*_ ∈ ℝ denote the site-specific effect for methylation site *g* and let *γ*_*ig*_ ∈ ℝ and *δ*_*ig*_ ∈ ℝ denote the additive and multiplicative effects for batch *i*, respectively. ComBat then assumes the following model:

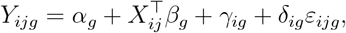

where 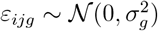 denotes the error term with standard deviation *σ*_*g*_ *>* 0. The model estimation and batch effect correction were performed in the following three steps:

### Step 1: Estimation of the Global Parameters

Global parameters *α*_*g*_, *β*_*g*_, and *σ*_*g*_ for each methylation site *g* were estimated to standardize the observed data so that the mean and variance were equal across methylation sites. First, we considered a model that included only the additive batch effect.

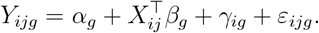

Let 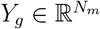 denote a vector of observations, defined as follows:

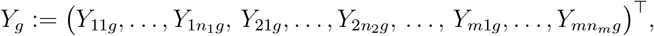

where 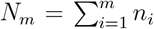 denotes the total number of samples. Let 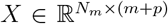 be the design matrix constructed from the batch indicator variables and other covariates, i.e., *X* := (*X*_batch_, *X*_cov_), where *X*_batch_ 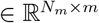 is the matrix of indicator variables for the *m* batches constructed without a constant column and 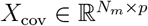 is the matrix of the *p* covariates defined as

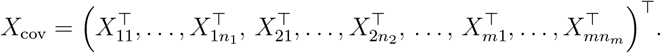

The estimates 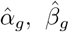, and 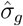 are then calculated under the identifiability condition 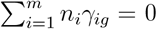. First, the ordinary least squares solution 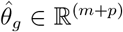 is calculated as

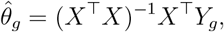

where 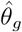 can be partitioned as:

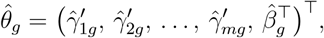

where 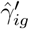 is the estimate of *γ*_*ig*_ without centralization. Then, 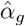 is calculated as:

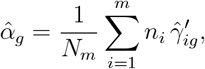

and 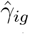 is 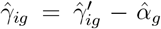 satisfying 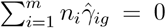. Furthermore, the variance 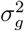 is estimated as:

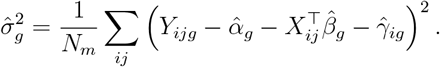

### Step 2: Estimation of the Batch Effect Parameters

Using the estimates of global parameters, the observed data were standardized as follows:

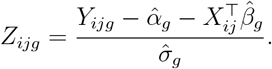

We assume the following Bayesian hierarchical model for the standardized data *Z*_*ijg*_:

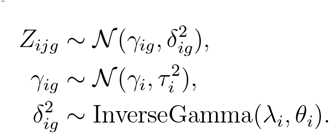

The hyperparameter estimates 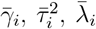, and 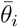 are computed using the method of moments, and are obtained as

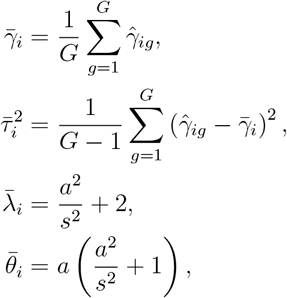

where 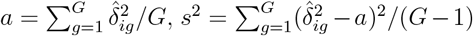, and 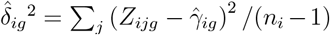. Then, the empirical Bayes estimators 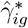 and 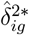 of batch effects satisfy the following simultaneous equations:

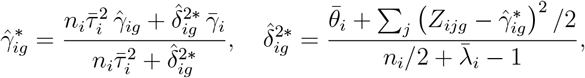

and were obtained as numerical solutions through iterative updates.

### Step 3: Data Correction

Finally, using the estimated global and batch effect parameters, the corrected data were obtained as follows:

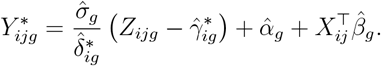

## 3 Proposed Incremental Framework

We propose Incremental ComBat (iComBat), an incremental framework based on ComBat that corrects a newly included batch, avoiding modification of the existing correction results. This section describes the proposed method in detail.

After applying ComBat to the existing batches *i* = 1, …, *m*, the estimated parameters for the correction model for each existing batch and for each site *g* = 1, …, *G*, namely 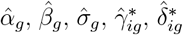, and the hyperparameter values of the prior distributions 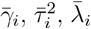, and 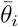, were obtained. For a newly added batch *i* = *m* + 1, we propose correcting the new data by estimating the batch effect parameters 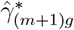 and 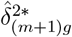 using the new batch and the global parameters estimated using the existing batches {1, …, *m*}. Specifically, to standardize the expression values *Y*_(*m*+1)*jg*_ in the new batch *m* + 1 at each site *g*, the estimated parameters 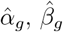, and 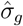 from existing batches were employed. That is,

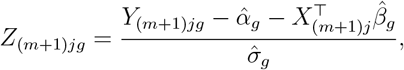

Using the standardized data *Z*_(*m*+1)*jg*_, the estimates 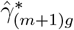 and 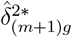 are obtained using the same procedure as in Step 2 of the conventional ComBat. The corrected data for the new batch are then computed as:

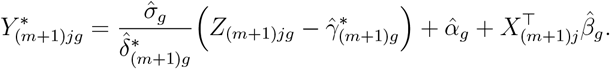

Subsequently, the corrected values 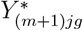 are obtained on the same scale as those of the existing batches.

## 4 Numerical Experiments

### 4.1 Settings

We evaluate the performance of the proposed method using numerical experiments. For methylation sites *g* ∈ {1, …, *G*} and samples *j* ∈ {1, …, *n*_*i*_}, data *Y*_*ijg*_ belonging to batch *i* ∈ {1, …, *m*} were generated according to the following model:

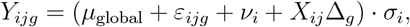

where *μ*_global_ is the global mean, 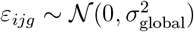 is the noise term with global variance 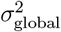, *ν*_*i*_ is the mean shift owing to batch *i*, and *σ*_*i*_ is the scaling factor for batch *i. X*_*ij*_ ∈ [0, 1] is an indicator variable representing the group to which sample *j* belongs (e.g., 0: the control group; 1: the treatment group). Δ_*g*_ denotes the group effect.

For the data generation, the total number of sites was set to *G* = 500. To introduce differential methylation between the control and treatment groups only for the first 50 sites, we set Δ_*g*_ = 0.5 for *g* = 1, …, 50, and Δ_*g*_ = 0 for *g* = 51, …, 500. The global parameters were set to *μ*_global_ = 5 and *σ*_global_ = 1. For the existing batches, we used *m* = 3 and set the following parameters: for batch 1, (*n*_1_, ν_1_, *σ*_1_) = (50, 0, 1); for batch 2, (*n*_2_, *ν*_2_, *σ*_2_) = (30, 2.5, 1.5); and for batch 3, (*n*_3_, *ν*_3_, *σ*_3_) = (40, −2, 1.2). Thus, the data from the existing batches comprised 120 samples. Furthermore, a new batch (*m* + 1 = 4) is added to the parameters (*n*_4_, *ν*_4_, *σ*_4_) = (30, −4, 1.8). Within each batch, the samples were equally assigned to the control and treatment groups.

iComBat was then applied to the generated data. The first three batches were corrected using the conventional ComBat, and the new batch was corrected using the global parameters obtained in the already applied ComBat. For comparison, simultaneous batch correction was performed for all four batches using ComBat. The experiment was repeated 1000 times. For uncorrected data, corrected data using ComBat, and corrected data using iComBat, Welch’s t-test was performed for each methylation site to compare the treatment and control groups. Using a significance level of 5%, the true-positive rate (TPR) and false-positive rate (FPR) were calculated. In addition, principal component analysis (PCA) was applied to the combined data from the existing and new batches, and scatter plots were drawn using the first principal component (PC1) and the second (PC2).

### 4.2 Results

Table 1 shows the average TPR and FPR for differential methylation site detection from 1000 simulation experiments using each correction method. Although the uncorrected data yield an average TPR of 0.017, the data corrected using ComBat exhibit an average TPR of 0.836. Furthermore, the data corrected using iComBat achieved an average TPR of 0.855, which is similar to the results obtained from simultaneous correction using the ComBat for all four batches. The average FPR was nearly zero in all cases.

**Table 1:** Average true positive rate (TPR) and false positive rate (FPR) for differential methylation site detection in 1000 simulation experiments.

| Correction Method | Average TPR | Average FPR |
| --- | --- | --- |
| No Correction | 0.017 | 0.0 |
| ComBat | 0.836 | 0.039 |
| iComBat | 0.855 | 0.047 |

Figure 1 shows the PCA plots of the uncorrected data and the data corrected using each method. For uncorrected data, the data from each batch were separated. However, for the data corrected using either ComBat or iComBat, a new batch (batch 4) was mapped such that it overlapped with existing batches (batches 1, 2, and 3). Thus, iComBat performs batch correction for new data without altering the correction results of the existing batches, while preserving the systematic signal associated with the assigned treatment.

**Figure 1:**
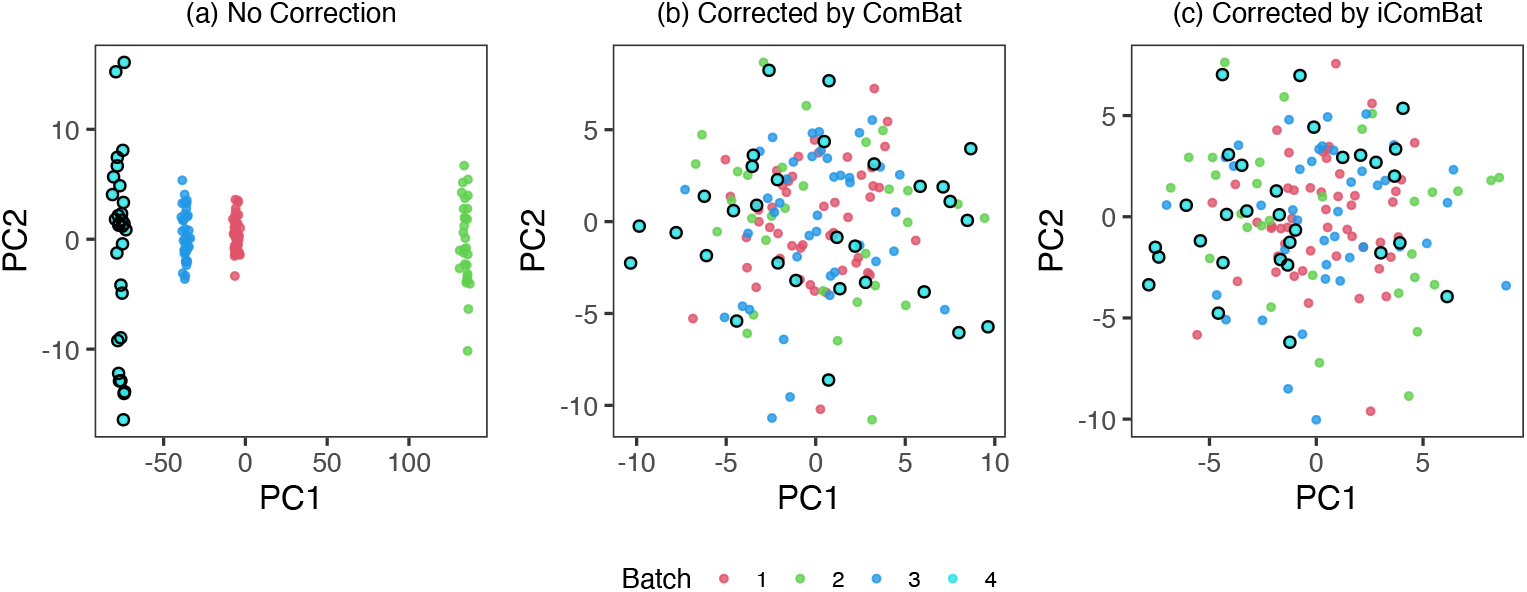
PCA plots of simulated data comprising original batches 1–3 and a newly added batch 4 (outlined in black): (a) Uncorrected data. (b) All four batches corrected together using standard ComBat. (c) Batches 1–3 corrected using ComBat, with batch 4 subsequently corrected using iComBat to align with the original batches.

## 5 Application to Actual Data

### 5.1 Settings

To demonstrate the practical performance of iComBat, we employed the publicly available GSE42861 dataset, which contains Illumina HumanMethylation450 measurements from whole-blood DNA of healthy controls and patients with rheumatoid arthritis [27]. Only samples annotated as “Normal” were retained, resulting in *n* = 166 control samples collected from 20 Sentrix array identifiers (6–11 samples per array; see Table 2). Each array was used as an independent batch. The raw beta values were transformed into M-values. Probes with zero variance within at least one batch were excluded. Then, there were *G* = 485, 577 CpG sites common to all samples.

**Table 2:** Sample counts for existing and new Sentrix arrays used in the incremental experiment. Sentrix-array identifiers are described as Batch ID.

| Existing |  | New |  |
| --- | --- | --- | --- |
| Batch ID | Count | Batch ID | Count |
| 5730192015 | 9 | 5730192007 | 7 |
| 5730192033 | 7 | 5730192034 | 8 |
| 5730053006 | 8 | 5730053009 | 11 |
| 5730053010 | 9 | 5730053012 | 8 |
| 5730053011 | 10 | 5730053030 | 6 |
| 5730053027 | 9 | 5730053039 | 8 |
| 5730053037 | 8 | 5730053040 | 7 |
| 5730053038 | 8 | 5730053041 | 11 |
| 5730053047 | 8 | 5730053046 | 8 |
| 5730053055 | 6 | 5730053048 | 10 |
| Total: |  | 166 samples |  |

We compared the following three scenarios: (i) no correction: raw data of 20 batches; (ii) Corrected using ComBat: all 20 batches corrected using the standard ComBat simultaneously; (iii) corrected using iComBat: the first ten batches (a total 85 samples) were designated as the existing batches and corrected using ComBat; each of the remaining ten batches (a total 81 samples) were corrected using iComBat and global parameter values from the initial ComBat. The results are shown as PCA plots based on 10, 000 CpGs with the largest variance.

### 5.2 Results

Figure 2 shows scatter plots for each correction scenario. Samples are colored according to their batch, and the ten new batches corrected incrementally are highlighted with black outlines. The uncorrected data exhibited batch-specific clusters along PC1 and PC2. Both ComBat and iComBat reduced the between-batch dispersion. In particular, the data corrected using ComBat and iComBat exhibited similar distributions. This result demonstrates that the proposed incremental framework performs well on real datasets.

**Figure 2:**
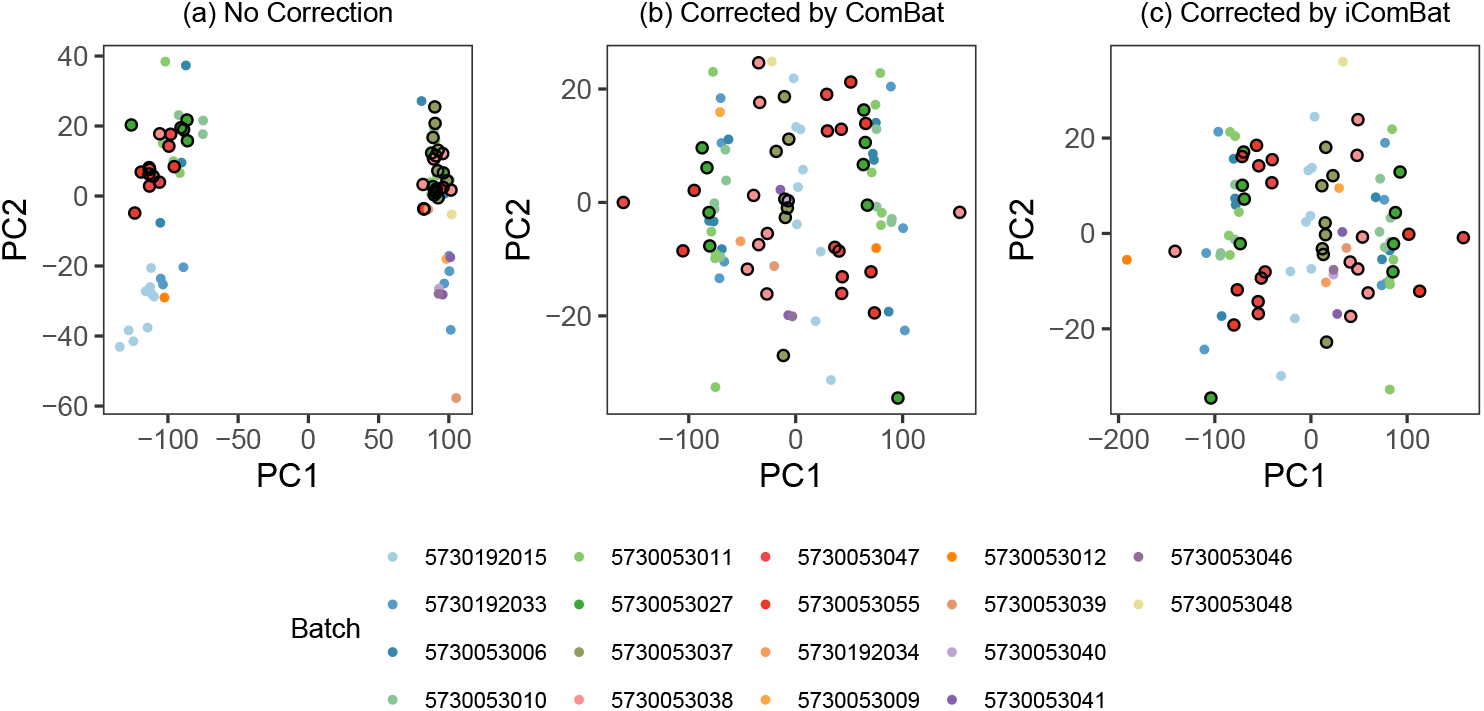
PCA plots of control samples from GSE42861 based on M-values: (a) Uncorrected data of 20 batches. (b) Data corrected using standard ComBat and the original 20 batches. (c) Additional ten batches processed with iComBat (black outlines) align with the ComBat-corrected ten batches.

## 6 Discussion

In this study, we extended ComBat, a widely used method for batch-effect correction of DNA methylation array data, by developing an incremental framework for batch effect correction [20]. The proposed method allows for the correction of new data without reprocessing existing, corrected data. Our numerical evaluations demonstrated that iComBat can achieve batch correction on new data without modifying existing batches, while preserving the power to detect systematic variations associated with treatment. The illustration using actual data of healthy controls in an EWAS study dataset showed that our method worked well on real data [27].

The proposed method is expected to be valuable for longitudinal studies involving repeated sample collection and measurements. For example, consider a clinical trial of an anti-aging intervention in which a change in the epigenetic clock is the primary endpoint [28]. When the baseline epigenetic clock value is incorporated into the inclusion/exclusion criteria or used for stratified randomization, it is necessary to evaluate the epigenetic clock values from baseline methylation data. To evaluate the change in the score, the epigenetic clock value was calculated from the methylation data at the end of the trial. However, if a batch effect correction scheme which differs from that applied at the baseline is used, the measured change may be biased. By employing iComBat, the baseline correction results can be preserved, while consistently correcting data at the end of the trial. This may have led to a more accurate evaluation of the intervention effects.

There are some variations in ComBat that may be extended to a similar incremental framework. For instance, methods have been proposed in the ComBat framework, which assumes that data follow a negative binomial or beta distribution [22, 23]. Their incremental versions may be formulated by following similar principles. Additionally, iComBat can be combined with other batch-effect correction methods. Comparative studies using large-scale longitudinal data have reported that a combination of quantile normalization and ComBat is effective in removing batch effects [29]. A similar preprocessing strategy combined with iComBat may yield more stable results.

## Acknowledgements

We would like to thank Editage (www.editage.jp) for the English language editing.

## Code Availability

The R implementation of iComBat is available from https://github.com/t-yui/iComBat.

## Data Availability

The raw DNA methylation data used in this study are publicly available from the Gene Expression Omnibus under accession GSE42861.

## Disclosure Statement

YT served as a technical advisor in statistical science for Rhelixa Inc., from April 2021 to March 2024. RN is the founder and chief executive officer of the company.

## Notes

https://github.com/t-yui/iComBat

https://www.ncbi.nlm.nih.gov/geo/query/acc.cgi?acc=gse42861

